# Phagocytosis in macrophages is regulated by the ATP-binding cassette family gene ABCF1

**DOI:** 10.1101/2023.09.05.556419

**Authors:** Hitesh Arora, Kyung Bok Choi, Lonna Munro, Cheryl G. Pfeifer, Wilfred A. Jefferies

## Abstract

Phagocytosis is a conserved biological mechanism that is integral to tissue remodeling, clearance of apoptotic cells, and immune defense in animals. Additionally, it serves as a pivotal means of sustenance for diverse unicellular eukaryotes. In the context of mammals, this crucial role is fulfilled by specific cell types such as macrophages, monocytes, dendritic cells, and neutrophils. It is orchestrated by an array of receptors, kinases, cytoskeletal elements, and enzymes, working collaboratively to enable the recognition, engulfment, and internalization of particles. Despite its profound significance, the intricate mechanisms underpinning the regulation of this phenomenon remains enigmatic. In this study, we present compelling evidence indicating the involvement of ABCF1, a member of the ATP-binding cassette family, in the FcγRIIA phagocytic pathway. ABCF1’s contribution lies in facilitating downstream signal activation through interactions with Src family members and SYK, pivotal players in this cascade. Additionally, our findings highlight ABCF1’s essentiality in the biosynthesis of various SFKs (Src family kinases) and MAPKs (mitogen-activated protein kinases). These molecules collectively oversee the orchestration of phagocytic cup formation, a pivotal step governing the engulfment process within macrophages. Consequently, the regulation of ABCF1 presents a potential avenue for modulating phagocytosis, allowing for the precise modulation of this fundamental process to either enhance or attenuate according to the specific physiological demands.

## Introduction

Cells have evolved many strategies to internalize particles from the extracellular space, namely pinocytosis, which refers to the uptake of fluid and solutes, receptor-mediated endocytosis, a process by which viruses and macro molecules enter the cell and lastly phagocytosis, which leads to actin dependent uptake of larger particles (>0.5 μm), like a pathogen or an apoptotic cell [1, 2]. Phagocytosis by macrophages can be mediated by many receptors namely Fcγ receptors (FcγR), Integrins, PRR, Apoptotic corpse receptors and Scavenger receptors [3]. Three classes of FcγR’s have been identified, namely FcγRI (CD64), FcγRII (CD36) and FcγRIII (CD16), each class containing several different isoforms [4]. Out of all the isoforms FcγRIIB is known to negatively regulate phagocytosis [5] and CD16 has been found to be a low affinity receptor, with one of its isoforms FcγRIIIA being expressed in macrophages at low levels and its other isoform FcγRIIIB is exclusively expressed in neutrophils [4]. Phagocytosis by FcγR’s is initiated by phosphorylation of tyrosine residues in the immuno-receptor tyrosine based activation motifs (ITAM) located in their cytoplasmic domain [6]. Activation and phosphorylation of ITAM has been attributed to SRC family kinases (SFK’s), which include Src, Lyn, Lck, Fyn, Yes, Fgr, Hck and Syk. Phosphorylation of ITAM’s by SFK’s leads to recruitment and activation of Syk. Syk in turn phosphorylates downstream adapter molecules like Src homology 2 domain containing leukocyte phosphoprotein of 76 KDa (SLP-76) [7]. Macrophages isolated from Hck-/-, Fgr-/-, Lyn-/- mice lead to reduced activation of Syk, thus reduced phagocytosis, but could still account for residual phagocytosis because these macrophages still expressed Yes, Src and Fyn kinases [8]. SFK’s have either been known to inhibit phagocytosis by phosphorylating inhibitory receptors containing immuno-receptor tyrosine based inhibitory motifs (ITIM’s) [9] or loss of SFK’s has been shown to impair actin polymerization below the phagocytic cup, and loss of Syk leads to failure to close the phagocytic vesicle [8]. Studies have also shown STAT3 dependent and MAPK independent activation of phagocytosis [10, 11]. After phagocytosis, the downstream response from these pathways leads to increase in anti-inflammatory cytokines and decrease in pro-inflammatory cytokines [11–13].

ABCF1 was the first mammalian ABC transporter found that lacked a transmembrane domain (TMD). The ABCF1 gene is located in the MHC class I region of the MHC locus on chromosome 6p21 in humans and on chromosome 17 in mice and is thought to be the mammalian homolog of the yeast protein GCN20. GCN20 is found to be associated with eIF2 and ribosomes through five different chromatography procedures and helps in translation initiation and its regulation by initiating production of GCN 4, protein that helps in translation up-regulation during amino acid deprivation [14, 15]. ABCF1, like GCN20 is known to be located in the cytoplasm and nucleoplasm, but not in the nucleolus [16]. It is also known to aid in eIF2 recruitment to 40S ribosomal subunit and initiate translation [17, 18]. It binds to ribosomes through the N-terminal domains and knockdown of ABCF1 by RNA interference impaired translation of both cap-dependent and -independent reporters GFP and CAT respectively [19, 20].

Unlike other ABC subfamilies A-D and G members, ABC subfamily E and F genes encode proteins do not function as transporters due to the lack of a TMD and are therefore, likely located in the cytoplasm and other cellular organelles. In our previously published work, we had created a heterozygous mouse model of another ABC gene, *Abcf1*+/-, after finding the homozygous knockout mouse was embryonic lethal at 3.5 days *post coitum* [21]. In a subsequent publication [22], we demonstrated that ABCF1 is a novel E2 ubiquitin conjugating enzyme that acts as a switch between the MyD88-directed pathway and the TRIF-TIRAM directed pathway of inflammation. These pathways either lead to a pro-inflammatory cytokine secretion or anti-inflammatory cytokine secretion and control macrophage polarization from M1 to M2 phenotype. By acting as a switch between these two pathways, ABCF1 subsequently protects against sepsis-mediated cytokine storm.

Here, in order to further explore the innate immune functions exerted by ABCF1, we make the discovery regulates essential phagocytosis mechanisms in macrophages.

## Results

### ABCF1 is necessary for anti-inflammatory cytokine and ISG production after 18 hours LPS stimulation

BMDMs, when stimulated with LPS, trigger a complex immune signaling with the production of both pro-inflammatory and anti-inflammatory cytokines. After LPS binds to TLR4, MyD88-dependent signaling is triggered as a result of an early phase response, followed by late phase TLR4 endocytosis and initiation of TRIF-dependent signaling [23]. To investigate if ABCF1 regulates LPS signaling, BMDMs were treated with *ABCF1* siRNA in presence and absence of LPS and levels of pro- and anti-inflammatory cytokines were analyzed. 20-30 fold increase in pro-inflammatory cytokines like IL-6, TNFα, IL-1β, IL-12p70, CXCL10, CXCL11 (**Figure 1A**) was observed when cells were treated with *ABCF1* siRNA and LPS and compared with LPS treated scrambled siRNA group. On the other hand, anti-inflammatory cytokines like IL-1Rα, IL-4, IL-10 and ISGs like IFNβ showed 30-fold downregulation with *ABCF1* siRNA treatment alone and the increase was not profound even after LPS stimulation (**Figure 1A**). Our data suggests that ABCF1 negatively regulates pro-inflammatory cytokine production after LPS stimulation and hence is necessary for anti-inflammatory cytokine production.

**Figure 1:**
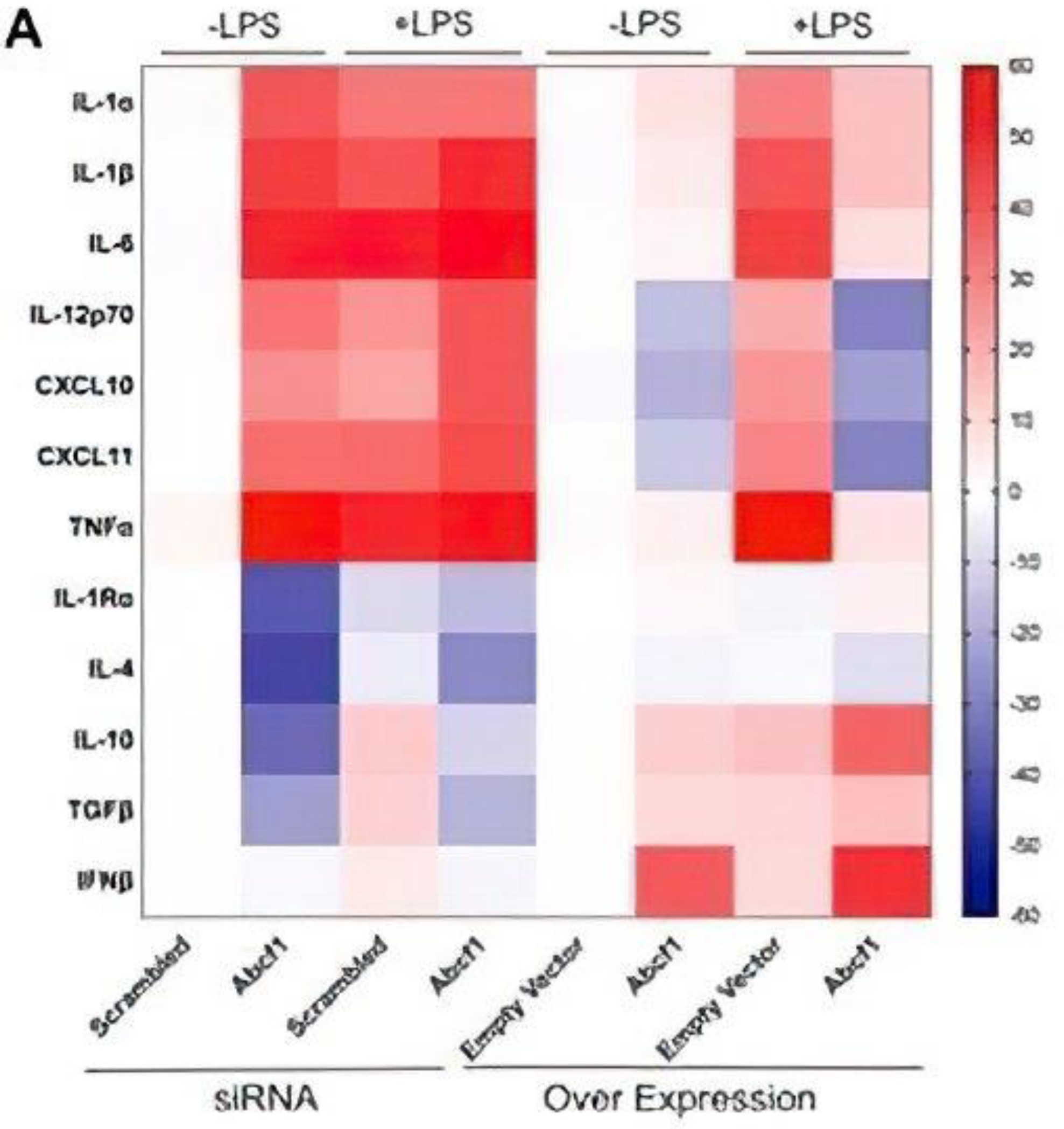
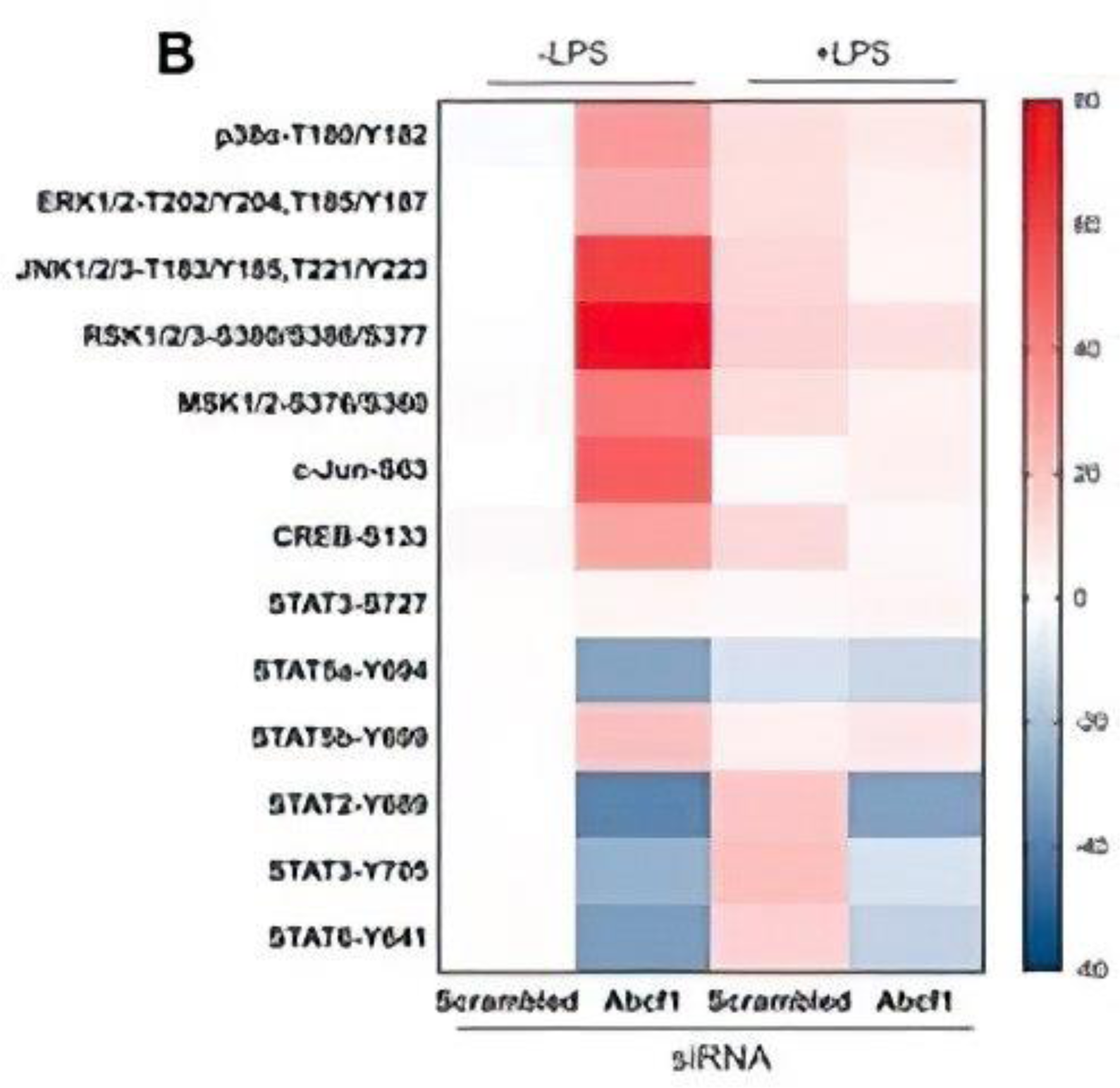
ABCF1 positively regulates anti-inflammatory cytokines and ISG specific transcription factors and negatively regulates MAPK after LPS stimulation: A) In bone marrow-derived macrophages (BMDMs), ABCF1 was manipulated through either overexpression (Over Exp.) or treatment with specific siRNA. These interventions were carried out both in the presence and absence of lipopolysaccharide (LPS) stimulation (at a concentration of 100 ng/ml for 24 hours). The resulting heat map illustrates the fold change in various cytokines and chemokines detected in the cell culture supernatants. B) Similarly, in BMDMs, ABCF1 was modulated using Abcf1-specific siRNA, and these cells were exposed to LPS treatment. The heat map presented here showcases the fold change in various Mitogen-Activated Protein Kinase (MAPK) and Interferon-Stimulated Gene (ISG)-specific phosphoproteins and transcription factors detected in whole-cell lysates (WCL).

Analysis of protein phosphorylation in BMDMs also showed that levels of MAPKs like p38, ERK and JNK were 40-50 fold elevated when treated with *ABCF1* siRNA and these levels significantly decreased when cells were stimulated with LPS (**Figure 1B**). MAPK specific transcription factor c-Jun also showed the same trend (**Figure 1B**). STAT transcription factors, on the other hand, showed an anomalous response. Phosphorylation levels of STAT 5a, STAT 2, STAT 3 and STAT 6 were downregulated after *Abcf1* siRNA and LPS stimulation (**Figure 1B**), whereas STAT 5b phosphorylation was negatively regulated with ABCF1 levels, when compared with LPS treated scrambled siRNA group.

### ABCF1 is necessary for FcγR IIA mediated phagocytosis in BMDM

Murine FcγRs have been shown to be critically involved in phagocytosis by macrophages [24, 25]. Out of the three murine FcγRs, FcγR I is a low binding receptor in macrophages [25], whereas the levels of FcγR I and II vary depending on the stimuli and cargo being engulfed by macrophages. We investigated the role of ABCF1 in regulating phagocytosis, by incubating *ABCF1* siRNA treated BMDMs in presence and absence of LPS with fluorescent labeled latex IgG coated beads. After flow cytometry analysis, 6 fold reduced fluorescent intensity was observed in *ABCF1* siRNA treated BMDMs when stimulated with LPS. This suggested that regulation by ABCF1 seems to be essential for phagocytosis of latex IgG coated beads after LPS stimulation (**Figure 2A**). Phagocytosis of fluorescent-labeled *E. coli* particles was also examined in *Abcf1* siRNA treated BMDMs. Absorbance value before and after BMDMs incubation with *E. coli* was measured and levels of phagocytosis was determined. After *E. coli* incubation, *ABCF1* siRNA treated BMDMs showed 4-fold reduced levels of phagocytosis, when compared with scrambled siRNA control; this suggested that ABCF1 seems to be necessary for phagocytosis of *E. coli* (**Figure 2A**). To investigate if ABCF1 mediated phagocytosis *via* which FcγR, levels of FcγRI (CD64), FcγRII (CD32) and FcγRIII (CD16) were analyzed via flow cytometry. Surface levels of CD64 and CD16 were not altered with loss of ABCF1 irrespective of LPS stimulation in BMDMs (**Figure 2B**). The levels of CD32 on the other hand are positively regulated by ABCF1 after LPS stimulation (**Figure 2C**). To analyze how ABCF1 mediates phagocytosis via CD32, the later was immunoprecipitated from ABCF1 overexpressed and LPS stimulated BMDMs. Strong protein bands were observed when the immunoprecipitates were immunoblotted with anti CD32 antibody (**Figure 2D**). The immunoprecipitates also showed strong protein bands when immunoblotted with anti phosphotyrosine antibody, suggesting CD32 ITAM phosphorylation in the presence of ABCF1, thus activating phagocytosis (**Figure 2D, 3**).

**Figure 2:**
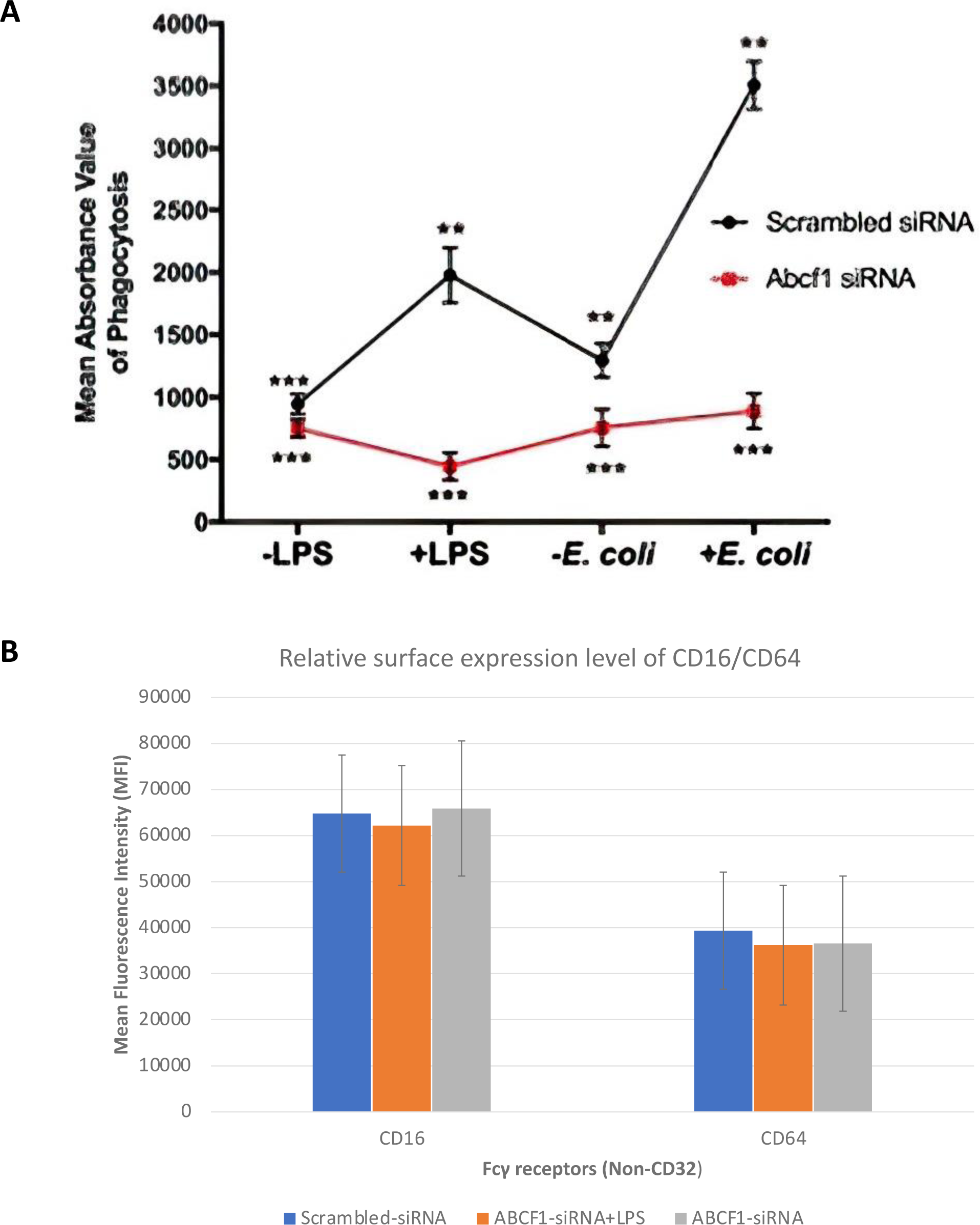
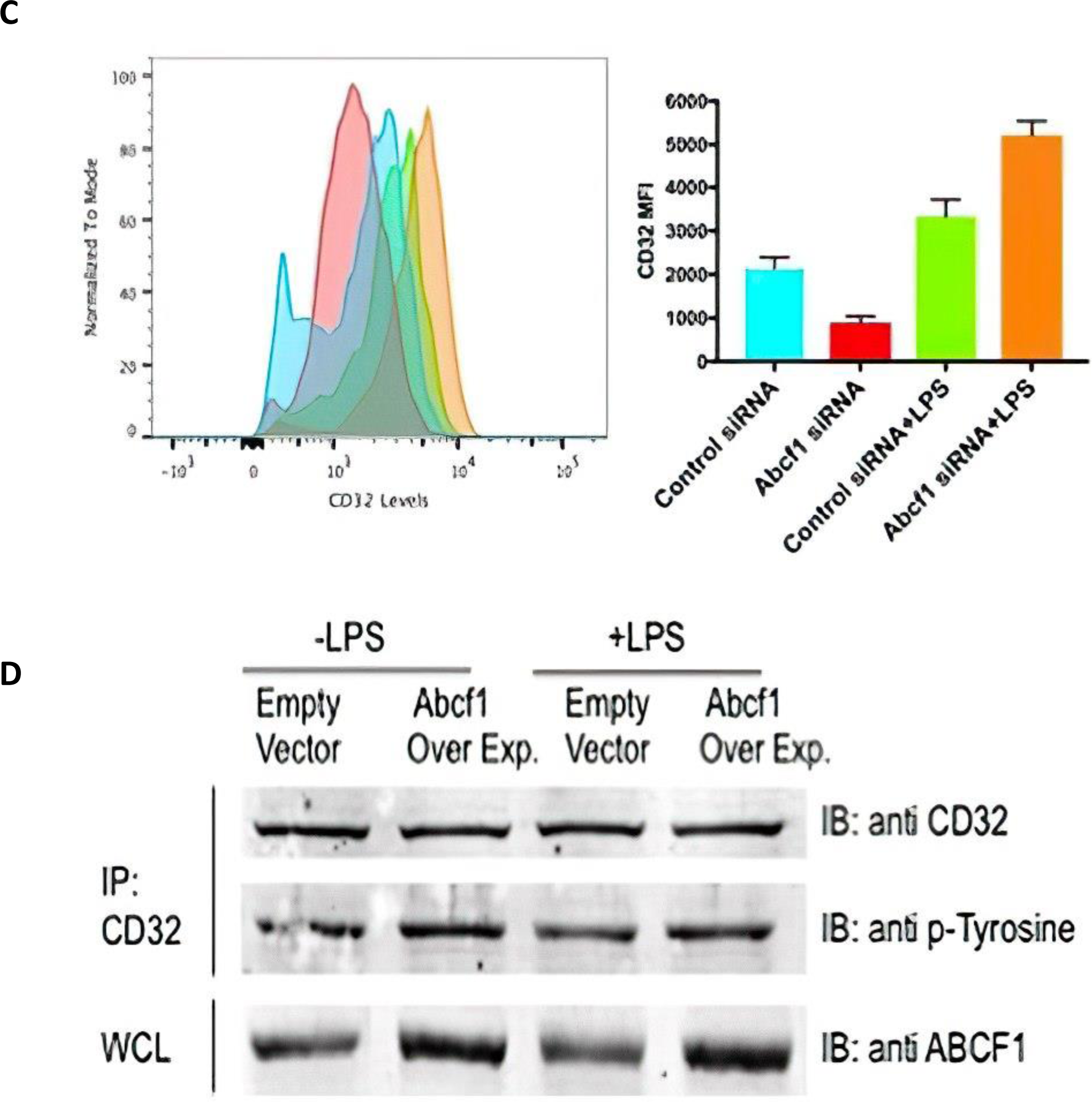
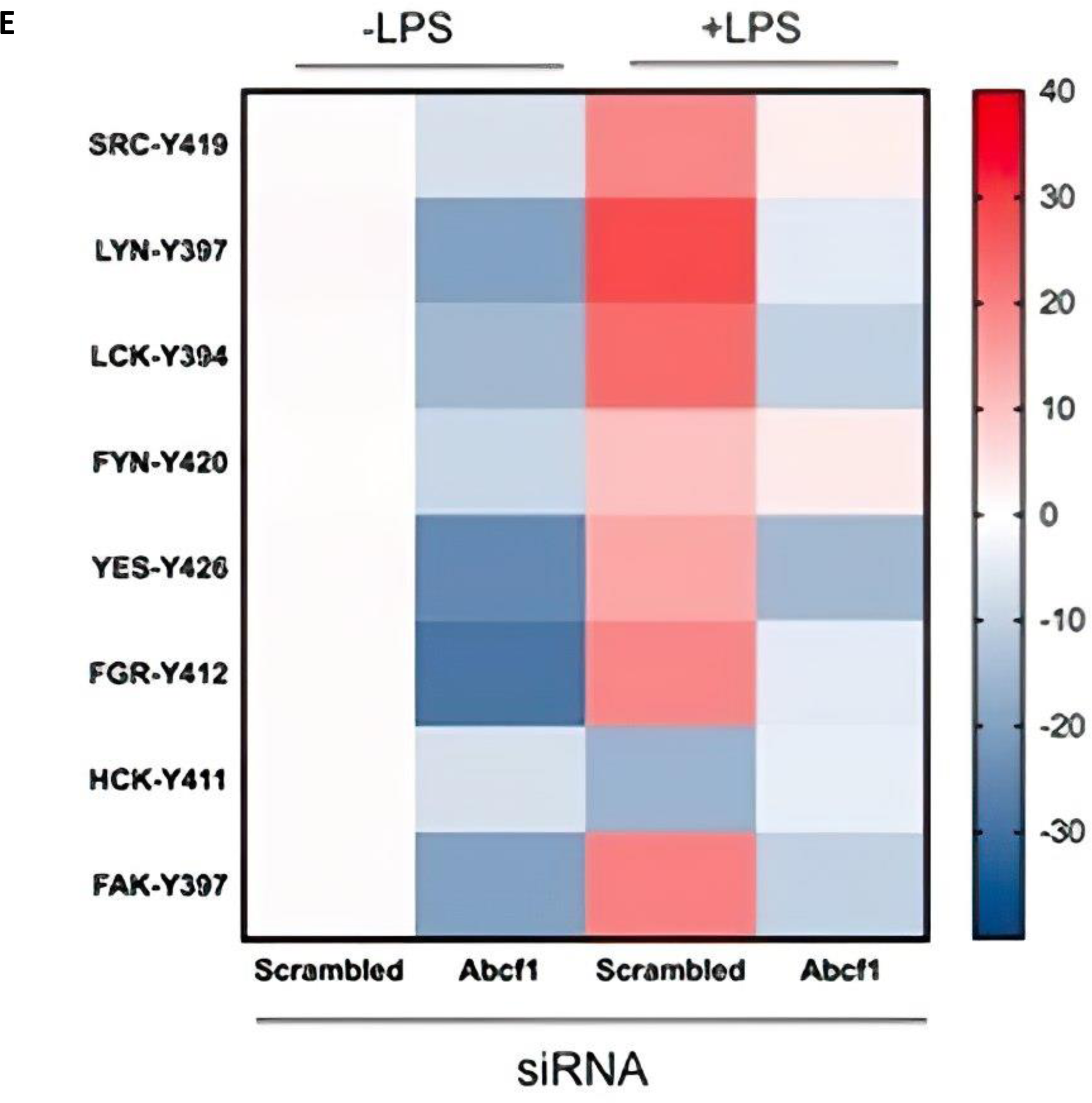
ABCF1 is necessary for FcgR II A mediated phagocytosis: **A)** The absorbance value associated with phagocytosis was assessed in bone marrow-derived macrophages (BMDMs) that were subjected to ABCF1-specific siRNA treatment. These BMDMs were incubated with fluorescently labeled IgG-coated latex beads for a duration of 2 hours, both in the presence and absence of lipopolysaccharide (LPS). Additionally, ABCF1-specific siRNA-treated BMDMs were incubated with fluorescently labeled E. coli for 2 hours. The results provide insights into the impact of ABCF1 modulation on phagocytosis; **B**) The levels of CD16 and CD64 were analyzed using flow cytometry in treated BMDMs. The bar graph depicts the change in mean fluorescence intensity (MFI), and error bars indicate the standard deviation. **C**) The levels of CD32 were analyzed using flow cytometry in treated BMDMs. The bar graph depicts the change in mean fluorescence intensity (MFI), and error bars indicate the standard deviation. This analysis sheds light on how ABCF1 manipulation influences CD32 expression; **C**) In BMDMs, ABCF1 was either overexpressed or not, and these cells were exposed to LPS. Whole-cell lysates (WCL) were immunoprecipitated with an anti-CD32 antibody. The immunoprecipitates were subsequently subjected to immunoblotting using anti-CD32 and anti-phosphotyrosine antibodies. This experimental approach allows the investigation of ABCF1’s role in CD32-associated signaling events; **D**) The heat map represents the fold change in phosphorylation levels of Src Family Kinases (SFKs) detected in whole-cell lysates (WCL) from ABCF1-specific siRNA-treated and LPS-stimulated BMDMs. This analysis provides insights into how ABCF1 modulation affects SFK phosphorylation levels in this context.

### ABCF1 positively regulates SFK and STAT3 activation and negatively regulates MAPK’s during phagocytosis

ITAM phosphorylation has been shown to recruit and phosphorylate SFKs, which triggers the downstream signaling cascade. SFKs like SRC, FYN, LYN and FGR, have been reported to be critically involved in phagocytosis [32, 33]. To investigate if ABCF1 regulates this mechanism, BMDMs were treated with *ABCF1* siRNA in presence and absence of LPS and phosphorylation levels of various SFKs were analyzed. Phosphorylation levels of all the SFKs were significantly downregulated with varying levels (Figure 2D), when cells were treated with *ABCF1* siRNA only. The levels increased negligibly in almost all SFKs with the exception of SRC and FYN, which were elevated 3-4 fold after LPS stimulation (**Figure 2E**).

## Discussion

Monocytes and macrophages are professional phagocytes that carry out the composite phenomenon of phagocytic engulfment. Phagocytosis is regulated by interplay of various cell-surface receptors and downstream proteins that carry out this signalling. Though phagocytosis has extensively been studied before, yet important gaps still remain in understanding its molecular pathway. We have shown that ABCF1 regulates FcγRII mediated phagocytosis (**Figure 2A,B**). ABCF1 seems to be essential for phosphorylation of tyrosine residue in the ITAM motif of FcγRII by SFKs, which leads to ubiquitination and subsequent phosphorylation of SYK and PLCγ2 (**Figure 2C**). This led to ABCF1 dependent phosphorylation of STAT transcription factors (**Figure 2D**), which regulate maturation of phagocytic cup and production of TRIF-dependent anti-inflammatory cytokines (**Figure 1A**).

Depending on the size of the cargo being engulfed, the mechanism of internalization of particles can be broken down into several distinct pathways, namely, clathrin mediated endocytosis (particle size <0.5 μm) and non-clathrin-mediated endocytosis, which includes phagocytosis and micropinocytosis (particle size >0.5 μm). Non-clathrin mediated micropinocytosis has recently been shown to be regulated by Epidermal Growth Factor Receptor (EGFR) and K-Ras [26], and loss of K-Ras regulates ABCF1 levels [27] and as ABCF1 regulates phagocytosis through Fc RII, it could perhaps also regulate micropinocytosis (summarized in **Figure 3**).

**Figure 3:**
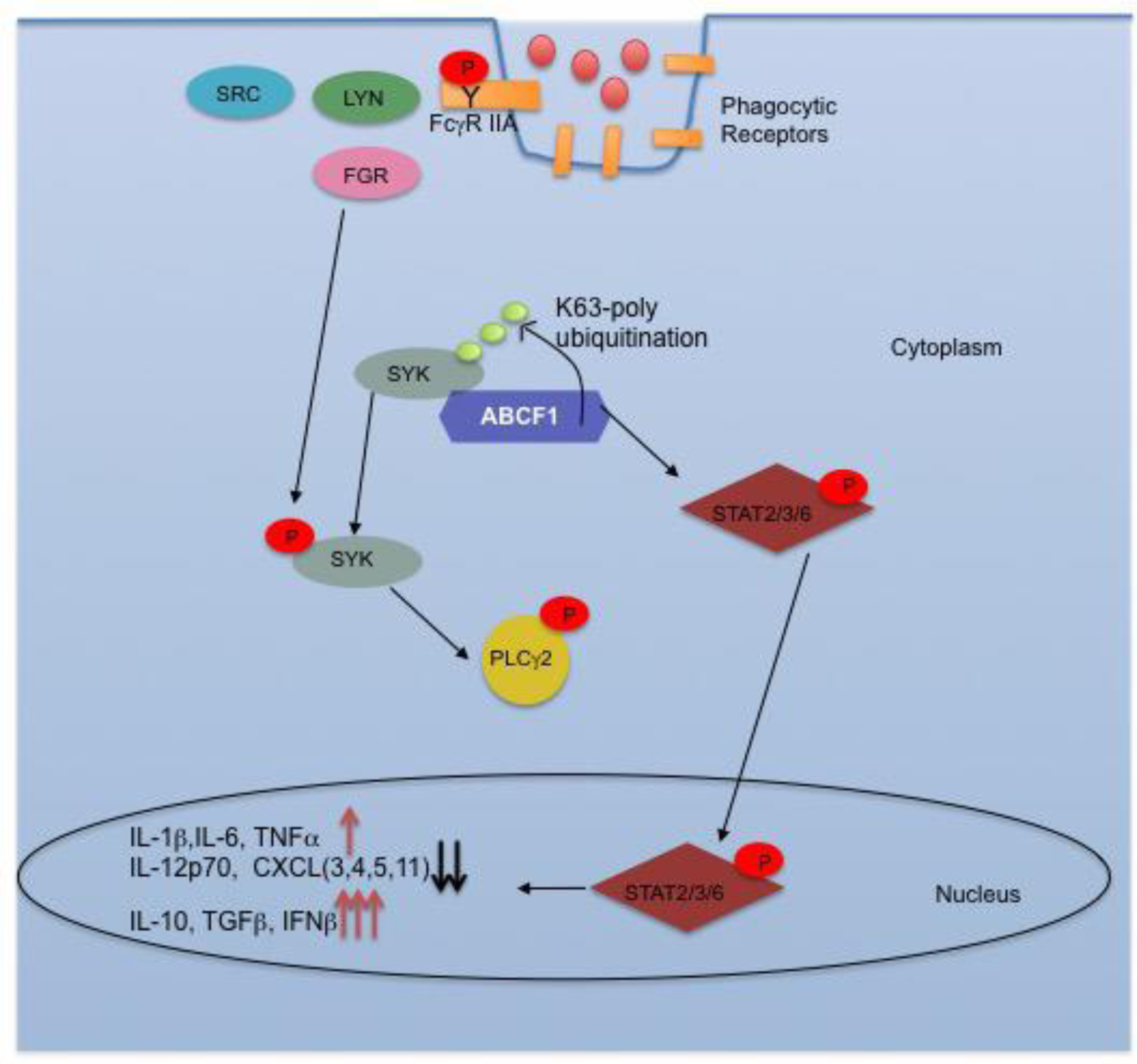
Model of ABCF1 regulation of Fc II A mediated phagocytosis: ABCF1 plays a crucial role in orchestrating the intricate process of phagocytosis, particularly in the engulfment of E. coli particles mediated by Fcγ receptors (FcγRs). The sequence of events involving ABCF1 in this process unfolds as follows: **1**) ABCF1 is essential for the phosphorylation of Immunoreceptor Tyrosine-Based Activation Motifs (ITAMs) by Src Family Kinases (SFKs) such as SRC, LYN, or FGR. This initial phosphorylation event is the trigger for FcγR-mediated phagocytosis of E. coli particles; **2**) Following ITAM phosphorylation, ABCF1 takes on the role of targeting Spleen Tyrosine Kinase (SYK) for K63-linked polyubiquitination. This ubiquitination event is crucial for subsequent signaling events; **3**) SYK, once polyubiquitinated, undergoes phosphorylation mediated by SFKs, such as FYN and other members of the SFK family. These phosphorylation events initiate downstream signaling pathways; **4**) Phosphorylated SYK sets off a cascade of events, including the phosphorylation of Phospholipase C Gamma 2 (PLCγ2), FYN, and other SFKs. These signaling molecules play critical roles in actin polymerization and the formation of the phagocytic cup, a structural prerequisite for successful phagocytosis; **5**) The downstream consequences of this signaling cascade include elevated phosphorylation levels of Signal Transducer and Activator of Transcription (STAT) proteins, particularly STAT 2, 3, and 6. These phosphorylated STAT proteins contribute to an increase in the production of anti-inflammatory cytokines. In summary, ABCF1’s involvement in phagocytosis of E. coli particles revolves around its role in initiating and coordinating a series of signaling events that ultimately lead to successful engulfment and the production of anti-inflammatory cytokines.

ABCF1 also regulates other innate immune functions. Previous studies have shown that a cell counteracts dsRNA infection via Oas1-RNase L mediated viral degradation [28–30]. Our studies also suggest that ABCF1 regulates this signaling pathway by interacting with OAS1a and controlling its 2’5’A activity. Immuno-precipitation experiments revealed that upon Poly I:C stimulation OAS1a was immuno precipitated with ABCF1. Protein analysis also showed that ABCF1 positively regulates OAS1a levels in *Abcf1* siRNA treated and Poly I:C stimulated BMDMs. ABCF1 was also found to negatively regulate protein levels of RNase L inhibitor, ABCE1, thus positively regulating RNase L dependent RNA degradation. It is noteworthy that its E2 ubiquitin enzyme activity does not appear to play a role in OAS1a regulation rather it mediates OAS1a phosphorylation. Thus, we provide evidence that ABCF1 positively regulate SFK and STAT transcription factors in LPS treated BMDMs, which aids in phagocytosis.

Based on these data a model for ABCF1 playing a pivotal role in meticulously orchestrating the intricate process of phagocytosis, specifically in the context of engulfing E. coli particles, a process mediated through Fcγ receptors (FcγRs) begins to emerge. This orchestrated sequence of events involving ABCF1 unfolds as follows: ITAM Phosphorylation: ABCF1 emerges as an indispensable factor in the initial phase of phagocytosis by facilitating the phosphorylation of Immunoreceptor Tyrosine-Based Activation Motifs (ITAMs). These ITAMs, located on Fcγ receptors, are phosphorylated by members of the Src Family Kinases (SFKs), including SRC, LYN, and FGR. This pivotal phosphorylation event serves as the ignition switch, setting in motion FcγR-mediated phagocytosis of E. coli particles. Ubiquitination of SYK: Following the successful phosphorylation of ITAMs, ABCF1 assumes a new role, directing its attention to Spleen Tyrosine Kinase (SYK). In a crucial step, ABCF1 targets SYK for K63-linked polyubiquitination. This ubiquitination process serves as a critical linchpin for the subsequent signaling cascade. SYK Phosphorylation: Once SYK undergoes polyubiquitination, it becomes primed for the next phase of signaling. Phosphorylation of SYK is the next domino to fall, and this process is catalyzed by SFKs, which include FYN and other members of the Src Family Kinases. These phosphorylation events initiate downstream signaling pathways, setting the stage for the ensuing cellular responses. Actin Polymerization and Phagocytic Cup Formation: Phosphorylated SYK acts as a signaling node that triggers a cascade of events. This includes the phosphorylation of Phospholipase C Gamma 2 (PLCγ2), FYN, and other SFKs. These signaling molecules play pivotal roles in actin polymerization, a dynamic process that is fundamental to the formation of the phagocytic cup. The phagocytic cup is a specialized cellular structure that envelops the target particle, preparing it for internalization. Anti-Inflammatory Cytokine Production: The downstream consequences of this orchestrated signaling cascade extend beyond the mechanical aspects of phagocytosis. Notably, the cascade culminates in elevated phosphorylation levels of Signal Transducer and Activator of Transcription (STAT) proteins, particularly STAT 2, 3, and 6. These phosphorylated STAT proteins act as key transcription factors that contribute to an upsurge in the production of anti-inflammatory cytokines. This shift in cytokine production represents an essential aspect of the host’s immune response, aiming to counteract inflammation and restore homeostasis. In essence, ABCF1’s pivotal role in the phagocytosis of E. coli particles revolves around its capacity to initiate and meticulously coordinate a sequence of signaling events (**summarized in Figure 3**). These events collectively ensure the successful engulfment of pathogens while simultaneously contributing to the production of anti-inflammatory molecules, a crucial facet of the immune response.

In summary, our findings provide substantial evidence that ABCF1 plays a positive regulatory role in Src Family Kinases (SFKs) and STAT transcription factors in LPS-treated BMDMs, thereby contributing to efficient phagocytosis and modulating downstream cellular functions. Given the involvement of multiple cell-surface receptors in phagocytosis, future research endeavors should focus on unraveling the intricate cooperation between ABCF1 and other receptors during phagocytosis and other related cellular functions. Nevertheless, ABCF1 presents an intriguing and novel entry point for exerting control over phagocytic processes, both in research and clinical applications.

## Materials and Methods

### Antibodies and Chemicals

#### Antibodies used for western blots and immunoprecipitation

1. Anti ABCF1 antibody (ThermoFisher Scientific; PA5-29955)
2. Anti ABCF1 antibody for co-immunoprecipitation (Proteintech; 13950-1-AP)
3. Anti CD16 antibody (BioLegend; S17014E)
4. Anti CD32 antibody (BioLegend; S17912B)
5. Anti CD64 antibody (BioLegend; X54-5/7.1)
6. Anti GFP antibody (abcam; ab290)
7. Anti GAPDH antibody (abcam; ab181602)
8. Anti Phsophotyrosine (abcam; ab10321)

#### Antibodies for Fluorescence Associated Cell Sorting (FACS)

1. CD11b (BioLegend; 101223).
2. CD86 (BioLegend; 105011).
3. CD206 (BioLegend; 1417097).
4. MHC-II (BioLegend; 141607).
5. TLR4 (BioLegend; 145406).

#### Chemicals used

1. Aprotinin (Santa Cruz Biotech; sc-3595).
2. Dynamin inhibitor I Dynasore (Santa Cruz Biotech; sc-202592).
3. LipoPolysaccharide (Santa Cruz Biotech; sc-221855).
4. N-Ethylmaleimide (Santa Cruz Biotech; sc-202719).
5. NP-40 (abcam; ab142227).
6. PMSF solution (Santa Cruz Biotech; sc-482875).
7. Smac Mimetic (Tocris; 5141).

### Cell and Animal Model Preparation

Bone marrow cells were harvested from the femur and tibia of C57BL/6 mice and subsequently differentiated into macrophages following the protocol detailed in reference 54. Briefly, bone marrow was collected from C57BL/6 mice by flushing their tibia and femur with Roswell Park Memorial Institute medium (RPMI) enriched with 10% heat-inactivated Fetal Bovine Serum (FBS). These bone marrow cells were then cultivated in 10 ml of RPMI supplemented with 10% FBS, glutamine, and 30% L929 cell supernatant containing macrophage colony-stimulating factor. The initial cell density was set at 10^6 cells/ml, and the cultures were maintained in 100-mm Petri dishes at 37°C with 5% CO2 in a humidified environment for 6 days. Subsequently, cells were collected using cold PBS, subjected to washing, resuspended in RPMI supplemented with 10% FBS, and utilized at a concentration of 2 × 106 cells/ml. All mouse experiments were approved by the Animal Care Committee at UBC.

### Transfection for Gene Knockdown and Overexpression

Gene knockdowns in cells were achieved through RNA interference (RNAi) using Lipofectamine™ RNAiMAX Transfection Reagent (ThermoFisher Scientific, catalog number 13778075) following the manufacturer’s instructions. The efficacy of knockdown was assessed via Western blot analysis. Specifically, ABCF1 gene knockdown was accomplished using Mouse Abcf1 siRNA from Santa Cruz Biotechnology (catalog number sc-140760), resulting in a consistent reduction of 79-81% compared to control samples treated with scrambled siRNA. For other genes, including mouse cIAP2 and mouse Trif, siRNAs from Santa Cruz Biotechnology (catalog numbers sc-29851 and sc-106845, respectively) were used for knockdown. Complementary gene over-expression studies were conducted using Lipofectamine™ 2000 Transfection Reagent (ThermoFisher Scientific, catalog number 11668027) following the manufacturer’s guidelines. The success of overexpression was verified through Western blot analysis.

### Phagocytosis Assay

The Phagocytosis Assay Kit (Green E. coli) (ab235900) (abcam, Canada) was employed that uses heat-killed, fluorescently pre-labeled E. coli particles for the precise detection and quantitative measurement of in vitro phagocytosis. This was accomplished using a spectrophotometer. Together with ABCF1 siRNA knockdown, the assay was used to measure either enhance or inhibit phagocytosis. The assay was performed as described by the [31].

### Phospho-kinase Analysis

Cell lysates were obtained from 2 × 10^7^ cells using the procedure outlined previously. These lysates were then applied onto nitrocellulose membranes featuring duplicated arrays of antibodies. These arrays were part of the Proteome Profiler Phospho-Kinase Array Kit, which is compatible with mouse samples and has been validated in previous studies (references 55 and 56). The analysis of phospho-kinase levels followed the manufacturer’s instructions, and the results were graphically depicted using bar graphs as detailed in the respective section.

### Cytokine Expression Analysis

The levels of cytokines were analyzed by preparing cell culture supernatant from 2 × 107 cells or blood serum. These samples were incubated on nitrocellulose membranes featuring duplicate arrays of anti-cytokine antibodies provided by the Proteome Profiler Mouse Cytokine Array Kit, Panel A, obtained from R&D Systems (catalog number ARY006). Treatments and incubations followed the respective experiment figure legends, and cytokine analysis was conducted in accordance with the manufacturer’s instructions. The resulting cytokine levels were quantified as fold changes and visually represented using bar graphs.

### Bar Graph Generation and Fold Change Calculation

Bar graphs were generated using GraphPad Prism software from San Diego, CA. To calculate the fold change of cytokine and phospho-kinase levels in Abcf1 siRNA-treated BMDMs (with or without PAMP treatment), we normalized the mean pixel density to scrambled siRNA-treated BMDMs (with or without PAMP treatment), respectively. Similarly, to determine the fold change of cytokine and phospho-kinase levels in scrambled siRNA- and PAMP-treated BMDMs, we normalized the mean pixel density to scrambled siRNA- and non-PAMP-treated BMDMs. This approach allowed us to quantify and compare the changes in cytokine and phospho-kinase levels under different experimental conditions.

### Western Blotting

Cell lysates were prepared as described above. For all co-immunoprecipitation (Co-IP) samples, the sample buffer was prepared without dithiothreitol (DTT), unless stated otherwise. In contrast, DTT was included in the sample buffer for all other samples intended for Western blot analysis. Electrophoresis and immunoblotting were performed using the specified antibodies. Western blot experiments were conducted in triplicate to ensure consistency and reliability.

### Statistical Analysis

Data are presented as the mean ± standard deviation (SD). Statistical comparisons between means were performed using paired or unpaired Student t-tests, as appropriate. A significance level of P < 0.05 was considered statistically significant. The sample size for each experiment is specified in the figure legend, indicating the number of observations or data points used in the analysis.

## Acknowledgements

We would like to thank Drs. Giorgia Caspani and Eliana Al Haddad for comments on the manuscript. This work was funded by operating grants to WAJ by the Canadian Institutes of Health Research (CIHR): MOP-86739 and MOP-133634; and by a Collaborative Research Agreement to the University of British Columbia by Mynd Life Sciences as well as by private donations to the <u>Sullivan Urology Foundation at Vancouver General Hospital</u> (https://www.urologyfoundation.ca). H.A. was supported by a Centre for Blood Research studentship. The funding sources had no role in the study design, data collection, analysis or interpretation of data, or in the writing of the paper.

## Author Contributions

Conceived Project: WAJ

Designed research: HA, CGP, WAJ

Performed research: HA, LM, KC

Analyzed data: HA, LM, CGP, KC, WAJ

Wrote paper: HA, WAJ

Edited paper: CGP, KC, WAJ

## Competing Interestss

All authors are equity holders in CAVA Healthcare Inc. the previous holder of IP related to this work subsequently acquired by Mynd Life Sciences Inc. and WAJ is a founder and is an equity holder in Mynd Life Sciences Inc, the current holder of UBC licenses and patents related to this work.

## Materials and Correspondence

Correspondence and material requests should be addressed to WAJ.

